# Enabling wider access to human molecular neuroscience research in pain: A simple preservation method for human dorsal root ganglion neurons in Hibernate A media

**DOI:** 10.1101/2025.09.06.674648

**Authors:** Joseph B. Lesnak, Mandee K. Schaub, Kimberly Gomez, Aida Calderon-Rivera, Santiago Loya-Lopez, Robert Stewart, Sooyeon Jo, Akie Fujita, Tomás Osorno, Marisa Desai, Keerthana Natarajan, Morgan K. Schackmuth, Marisol Mancilla Moreno, Stephanie I. Shiers, Anna Cervantes, Geoffrey Funk, Peter Horton, Erin Vines, Muhammad Saad Yousuf, Katelyn E. Sadler, Bruce P. Bean, Rajesh Khanna, Gregory Dussor, Theodore J. Price

**Affiliations:** Department of Neuroscience, Center for Advanced Pain Studies, The University of Texas at Dallas, Richardson TX, USA; Department of Pharmacology and Therapeutics, Center for Advanced Pain Therapeutics and Research (CAPToR), College of Medicine, University of Florida, Gainesville, FL, USA; Department of Neurobiology, Harvard Medical School, Boston, MA, USA; Southwest Transplant Alliance, Dallas, TX, USA

## Abstract

The use of human dorsal root ganglion (DRG) from organ donors opens the door for research into the molecular biology and physiology of human nociceptors; however, there are barriers to working with this tissue including logistical difficulties and limited access. We present an approach using Hibernate media to temporarily store either whole DRGs or dissociated DRG neurons prior to culturing and functional testing. Dissociation of DRGs following temporary storage (4-16hrs) in Hibernate media resulted in similar neuronal and immune cell yield as acutely dissociated DRGs. Neurons derived from DRGs stored in Hibernate media prior to dissociation exhibited similar electrophysiological properties and capsaicin responses as acutely dissociated DRG neurons. Similarly, neurons from acutely dissociated DRGs stored in Hibernate media (>24hrs) and shipped to geographically distant laboratories produced neuronal cultures displaying comparable electrophysiological properties as acutely cultured neurons. This approach overcomes insurmountable logistical burdens and increases access to freshly recovered human DRGs.

## Introduction

Dorsal root ganglion (DRG) neurons are responsible for detection of touch, temperature, proprioception, and nociceptive stimuli. These neurons play a key role in painful diseases where they become sensitized or develop spontaneous activity [8; 11]. Most research in this area has used rodent animal models usually relying on *in vivo* or *in vitro* physiological and pharmacological experiments using DRG cells, with an emphasis on DRG neurons. There is an increasing appreciation of species differences in DRG neurons that are likely important for translational research aimed at developing new therapeutics for pain [2; 14; 16; 18; 19]. Moreover, technological advancements have brought new priority to understanding the molecular composition of the human nervous system at the single cell level [2; 19; 23]. Investigators conducting animal model research schedule and conduct the experiments in their laboratories or in nearby facilities. Human tissue research that relies on tissue recovery from organ donors or rare surgeries is not scheduled by the investigators and requires travel to sites of recovery which are often not adjacent to research laboratories. Moreover, organ donor recoveries often occur during night hours when operating rooms are available in hospitals. Such logistical issues limit the availability of these tissues and put strain on the scientists doing the experiments, in particular when they disrupt sleep schedules and constrain geographic availability of tissues for research. We sought to address these issues by creating a human DRG tissue preservation protocol that can allow for more flexibility in scheduling and shipment of these precious human tissues for research purposes.

Neuronal tissue has been preserved in the past through the use of Hibernate media. This media was developed to preserve tissue in refrigerated temperatures and ambient CO_2_ for prolonged periods to aid in the storing and transporting of neuronal tissue [4]. When stored in Hibernate media supplemented with B27, rodent hippocampal tissue produces viable neuronal cultures even after 4 weeks of storage at 4°C [4]. Similarly, Hibernate media has been used to preserve cultures of Schwann cells [7] and oligodendrocyte precursor cells [22] at 4°C. Hibernate media has also been used as a transportation media for human cortical [3] and DRG tissue [10] between operating rooms and laboratories. Lastly, we have recently shown that human DRGs stored in Hibernate media prior to dissociation results in healthy immune cell isolation via fluorescently activated cell sorting [1]. However, it is unknown whether neurons isolated from human DRGs are altered following storage in Hibernate media. To directly address this gap in knowledge, we compared electrophysiological properties and responses to capsaicin of cultured human DRG neurons that were either 1) dissociated acutely, 2) stored as whole pieces of tissue in Hibernate media prior to dissociation, or 3) acutely dissociated then stored in Hibernate media and shipped to other laboratories for further analysis. We present functional data demonstrating that cultured human DRG neurons present similarly regardless of whether they were dissected acutely, preserved in Hibernate media prior to dissociation, or dissociated and stored in Hibernate media for shipment. This facile protocol has the potential to enable greater access to human DRG tissues and should allow greater flexibility for researchers working with organ procurement organizations on DRG recoveries for research purposes.

## Methods

### Human DRG Recovery

Lumbar and thoracic DRGs were recovered from human organ donors through a collaboration with the Southwest Transplant Alliance in Dallas, TX. Human DRGs were surgically recovered as previously reported within 4 hours of cross-clamp [17; 21]. In the operating room, DRGs were immediately placed in cold artificial cerebrospinal fluid [21; 24] (<1 hour) or BrainBits Hibernate A media without calcium (Fisher Scientific, NC0176976, Patent # WO2017123759A1) supplemented with 1% N2 Supplement-A (Stemcell Technologies, 07152), 2% NeuroCult SM1 (Stemcell technologies, 05711), 1% penicillin/streptomycin (Thermo Fisher Scientific, 15070063), 1% Glutamax (Thermo Scientific, 35050061), 2 mM Sodium Pyruvate (Gibco, 11360-070), and 0.1% Bovine Serum Albumin (BSA; Biopharm, 71-040) (4-16 hours) at 4 °C. DRGs were stored with at least 5 mL of Hibernate media per ganglion in a conical tube with a minimum of 10 mL per tube; 1–2 DRGs received 10 mL, and 3–4 DRGs received 20 mL. A total of 85 DRGs were used from 44 organ donors (N = 32 males, N = 12 females). Information on donor demographic and DRG levels used is provided in **Table 1**. All human tissue procurement procedures were approved by the University of Texas at Dallas Institutional Review Board following protocol Legacy-MR-15-237. DRGs were recovered by the Southwest Transplant Alliance who obtains informed consent for research tissue donation from a donor’s first-person consent (driver’s license or legally binding document) or legal next of kin. The United Network for Organ Sharing approves all policies on donor screening and consent. The Southwest Transplant Alliance follows the standards and procedures established by the US Centers for Disease Control and are inspected biannually by the Department of Health and Human Services. Distribution of any pertinent donor medical information follows HIPAA regulations to protect privacy.

**Table 1.** Donor Demographic and DRG Level Information.

### Human DRG Dissociation and Culturing

In all experiments, human DRGs were dissociated as previously reported [24]. Briefly, the bulb of the DRGs were exposed by trimming off excess connective tissue, fat, and nerve roots. The bulb of each DRG was diced into 3 mm x 3 mm sections and placed in 5 mL of prewarmed (37 °C) digestion enzyme consisting of 1 mg/mL of Stemxyme I (Worthington Biochemical, LS004106), 0.1 mg/mL of DNAse I (Worthington Biochemical, LS002139), and 10 ng/mL of recombinant human β-NGF (R&D Systems, 256-GF) in HBSS without calcium and magnesium (Thermo Scientific, 14170-112). The tubes were placed in a 37 °C shaking water bath with light trituration every hour until the DRGs were dissociated (3.5-13 hours). Samples were filtered through a 100 µm mesh strainer (Corning, 431752) and the resultant cell suspension was gently layered over 3 mL of 10% BSA in HBSS in a 15 mL tube. The tubes were centrifuged at 900 g for 5 minutes at room temperature. The supernatant was aspirated and the pellet resuspended in prewarmed DRG media (BrainPhys^®^ media (Stemcell technologies, 05790) containing 1% penicillin/streptomycin, 2% NeuroCult SM1, 1% Glutamax, 1% N2 Supplement-A, 10 ng/mL recombinant human β-NGF, 2% HyClone™ Fetal Bovine Serum (Thermo Fisher Scientific, SH3008803IR), and 0.1% of 3 mg/mL 5-Fluoro-2′-deoxyuridine,thymidylate synthase inhibitor (Sigma-Aldrich, F0503) and 7 mg/mL uridine (Sigma-Aldrich, U3003). The number of neurons yielded per DRG were estimated through manually counting the number of neurons in a 10 µl aliquot of the resulting cell suspension. Cells were plated on either 12 mm coverslips or 24 well glass bottom plates pre-coated with 0.1 mg/mL of poly-D-lysine (Sigma-Aldrich, P7405-5MG) depending on the experimental assay. Cells were incubated at 37°C and 5% CO_2_ for 3-4 hours to allow for adherence. Following adherence, wells were flooded with prewarmed media and half media changes were performed every other day.

In a separate set of experiments, the resultant neurons were preserved in Hibernate media instead of being plated immediately. In this case, following dissociation and BSA filtering as mentioned above, the resulting pellet was resuspended in 10 mL of pre-chilled Hibernate media and stored at 4 °C. The next day, the cells were shipped overnight at 4°C to either the laboratories of Dr. Rajesh Khanna at the University of Florida or Dr. Bruce Bean at Harvard University via priority overnight FedEX shipping. Cells were shipped in standard Styrofoam boxes with cold gel packs to maintain temperature close to 4°C during shipment. Between resuspension of neurons at UTD and arrival of tubes at each university, neurons were stored in Hibernate media at 4 °C for 16-42 hours.

### Fluorescently Activated Cell Sorting

For fluorescently activated cell sorting (FACS), DRGs were similarly dissected and dissociated in 5 mL of pre-warmed digestion enzyme consisting of 2 mg/mL of Stemxyme I, 0.1 mg/mL of DNAse I, and 10 ng/mL of recombinant human β-NGF in HBSS. The cell suspension was filtered through a 70 µm mesh strainer (CELLTREAT Scientific Products, 229483) and centrifuged (350 g, 5 minutes, room temperature; these settings were used for all subsequent centrifugation steps). The supernatant was removed, and the pellet was resuspended in 1X Red Blood Cell Lysis Buffer (Biolegend, 420301) and incubated for 5 minutes at room temperature. The cells were centrifuged, the supernatant was removed, and the pellet was resuspended in 0.5% BSA in 1X phosphate buffered saline (PBS). Cells were incubated with myelin removal beads (Miltenyi Biotec, 130-096-433) for 15 minutes at room temperature. The cells were washed with 1 mL of 0.5% BSA in 1X PBS, centrifuged, resuspended in 0.5% BSA in PBS, and filtered through a LS column (Miltenyi, 130-042-401) attached to the MidiMACS separator (Miltenyi, 130-042-301) according to the manufacturer’s recommended protocol. Cells were centrifuged, resuspended in 1X PBS, and proceeded to cellular staining.

Cells were first incubated with a Zombie UV live/dead stain (Biolegend, 423107) for 10 minutes at room temperature, protected from light. Cells were washed with 1 mL of flow cytometry staining buffer (Invitrogen, 00-4222-26), spun down, and resuspended in flow buffer. Cells were incubated with a human Fc receptor blocker (TruStain FcX, Biolegend, 422302) for 10 minutes at room temperature, protected from light. Cells were then incubated with antibodies targeting CD45, CD11b, and CD3 for 30 minutes on ice, protected from light (See **Supplemental Table 1** for antibodies used in the FACS experiments). Cells were washed with 1 mL of flow buffer, centrifuged, and resuspended in flow buffer and kept on ice until sorting. FACS was used to collect live, CD45+, and either CD11b+ or CD3+ cells on a BD FACSAria Fusion using a previously reported gating strategy [1].

### Immunocytochemistry

For immunocytochemistry experiments, human DRG cultures were seeded at a neuronal density of 50-100 neurons per well in glass bottom 24 well plates (Cellvis, P24-1.5H-N). On DIV 5, cells were washed once with 1X PBS and fixed with 4% paraformaldehyde (Electron Microscopy Sciences, 15710) in 1x PBS for 15 minutes at room temperature in the dark. Cells were then washed 3X with 1X PBS and blocked with 10% normal goat serum (R&D Systems, S13150H) in PBS for 1 hour at room temperature. Cells were permeabilized with 0.3% Triton X-100 (Sigma-Aldrich, X100) and 10% NGS in 1X PBS for 30 minutes at room temperature. Cells were then incubated with a primary antibody targeting peripherin (1:1000, Encor, CPCA-Peri) overnight at 4 °C. The next day cells were washed 3X with 1X PBS and incubated with goat anti-chicken 488 secondary antibody (1:2000, Invitrogen, A11039) and DAPI (1:5000, Cayman Chemical, 14285) for 1 hour at room temperature. Cells were then washed 3X with 1X PBS and wells were flooded with 500 µL of PBS and kept in the dark at 4 °C until imaging. Cells were imaged on an Olympus FV3000 confocal microscope at the University of Texas at Dallas.

### University of Texas at Dallas (UTD) - Electrophysiology

For electrophysiology experiments, DRG neurons were plated at ∼50 neurons per well on 12 mm coverslips in a 24 well plate. On the day of testing, DRG neurons were transferred from DRG media to bath solution (135 mM NaCl, 10 mM Glucose, 10 mM HEPES, 5 mM KCl, 2 mM CaCl_2_, 1 mM MgCl_2_, 20 mM sucrose, with a pH of 7.4 and osmolarity ranging from 310-320 mOsm/L) and all data was collected within 90 minutes of transfer. Whole-cell patch clamp electrophysiology was conducted using a Multiclamp 700B (Molecular Devices, San Jose, CA) amplifier and pClamp10 acquisition software (Molecular Devices, San Jose, CA). Glass micropipettes (outer diameter 1.5 mm: inner diameter, 1.10 mm; Sutter Instruments, BF150-110-10) were pulled using a PC-100 puller (Narishige) and fire polished to the resistance of 1-3 MΩ using a microforge (MF-83, Narishige). The pipettes were then filled with pipette solution (135 mM KCl, 10 mM HEPES, 4 mM ATP-Mg, 0.9 mM GTP-Na, 0.5 mM EGTA, 5 mM NaCl with a pH of 7.4 and osmolarity ranging from 305-320 mOsm/L) and pipette current was zeroed. The pipette was then attached to the cell and suction was applied until a giga-ohm seal was achieved. The membrane was ruptured to achieve whole-cell mode and then compensation for fast and slow capacitance was applied. A -70 mV to 0 mV to -70 mV voltage step was applied in voltage clamp to test for presence of voltage-activated inward and outward currents as an initial test for cell health. In current clamp mode, a 90 second recording with no stimulus was taken to assess spontaneous activity. Next, a series of 20-millisecnd current steps incrementing current with a delta of 10 pA was applied until the cell fired the first action potential to measure rheobase. Next, a series of 1-second ramp current injections was applied to the cell, starting with a ramp from 0-100 pA and increasing the ramp ending voltage by a delta of 200 pA up to maximum of 2900 pA. Finally, a series of 500-millisecond step current injections starting with 0 pA and incrementing by 50 pA up to a maximum of 2100 pA was injected to assess action potential firing as a function of current injection. Data analysis was conducted manually using Clampfit 10.4 software (Molecular Devices).

### University of Texas at Dallas - Calcium Imaging

For calcium imaging experiments, human DRG neurons were plated at 100-150 neurons per coverslip on 12 mm coverslips in a 24 well plate. On DIV 3-5, the coverslip was removed from the well containing DRG media and placed in a well with calcium imaging bath solution (145 mM NaCl, 3 mM KCl, 2.5 mM CaCl_2_, 1.2 mM MgCl_2_, 10 mM HEPES, 7 mM Glucose, pH 7.4±0.1, osmolarity 320±3 mOsm/L). The coverslip was then incubated with the calcium indicator Fura 2 (3 ug/mL; Thermo Fisher Scientific, F1221) in bath solution containing 2% BSA for 45 minutes at room temperature in the dark. The coverslip was then incubated in bath solution for 15 minutes at room temperature in the dark to allow for de-estertification of the Fura dye. Coverslips were mounted onto an inverted Nikon Eclipse Ti2 fluorescent microscope and images were captured at both 340 nm and 380 nm excitation. Cells were perfused with bath solution for 90 seconds, followed by stimulation with capsaicin (200, 20, or 2 nM; Sigma-Aldrich, M2028) for 2 minutes. Coverslips were perfused again with bath solution for 2 minutes and then treated with 50 mM KCl to confirm neuronal viability. The ratio of the 340/380nm fluorescent signal, with a background correction, was calculated for each neuron using NIS elements software (Nikon, Version 6.10.01). A neuron was considered a capsaicin responder to test stimuli if it exhibited a ≥25% increase in the 340/380 nm ratio from baseline. A neuron was considered viable and included in the analysis if it exhibited a ≥25% increase in 340/380 nm ratio following KCl stimulation.

### University of Florida – Electrophysiology

The Hibernate media cell suspension was centrifuged at 350 g for 3 minutes. The supernatant was removed and the cells were resuspended in BrainPhys media (STEMCELL Technologies, 05790) supplemented with 25 ng/ml beta-NGF (R&D systems, 256-GF-100) and 2.5 ng/ml GDNF (R&D systems, 212-GD-010) in addition to 1% GlutaMax, 1% N2, 2% SM1, and 1% penicillin/streptomycin. The cells were seeded on 35 mm ibiTreat plates (ibidi GmbH, 80136) and incubated at 37 °C and 5% CO_2_ with half media changes performed every 3 days. The electrical activity was assessed on DIV 2-3. For current-clamp recordings, the external solution contained 154 nM NaCl, 5.6 mM KCl, 2 mM CaCl_2_, 1 mM MgCl_2_, 10 mM D-Glucose, and 8 mM HEPES (pH 7.4 adjusted with KOH, and mOsm/L= 300). The internal solution comprised: 137 mM KCl, 10 mM NaCl, 1 mM MgCl_2_, 1 mM EGTA, and 10 mM HEPES (pH 7.3 adjusted with KOH, and mOsm/L= 277). Recordings of action potentials were made at room temperature in whole-cell patch clamp configuration and current-clamp mode. DRG neurons with a resting membrane potential (RMP) more hyperpolarized than −40 mV, stable baseline recordings, and evoked spikes that overshot 0 mV were used for experiments and analysis. Action potentials were evoked by a ramp pulse from 0-1000 pA in 1 second. Results were analyzed using Fitmaster software version 2x92 (HEKA) and Easy Electrophysiology 2.7.3.

### Harvard University – Electrophysiology

Cell were received in Hibernate media and were spun down at 620 RPM (65g) then resuspended in culture media that consisted of BrainPhys media (Stemcell technologies, #05790), 1% penicillin/streptomycin,1% GlutaMAX (Gibco, #35050-061), 2% NeuroCult SM1 (Stemcell technologies, #05711), 1% N-2 Supplement (Thermo Scientific, #17502048), 2% Fetal Bovine Serum (Gibco #A5256801). Cells were then plated in 24 well plates on 12 mm coverslips (Fisherbrand #12-545-80) that had previously been coated with 0.01% Poly-D-Lysine overnight at 4 °C. Poly-D-Lysine was removed and coverslips were allowed to dry before plating of cells. After plating of the cells, the plates were incubated at 37 °C for 1-2 hours so that cells would adhere to the coverslips before wells were flooded with 1 mL of culture media. Plates were then kept in in a 5% CO2 incubator set at 37°C for up to 7 days, with half of the media in each well exchanged every 2-3 days. Cell were used up to 7 days after plating. For recording, coverslips were placed into the recording chamber containing about 2 mL of Tyrode’s solution consisting of 155 mM NaCl, 3.5 mM KCl, 1.5 mM CaCl2, 1 mM MgCl2, 10 mM HEPES, 10 mM glucose and pH adjusted to 7.4 with ∼5 mM NaOH. Whole-cell patch clamp recordings were made using patch pipettes pulled from borosilicate capillaries (VWR International, 53432-921) using a Sutter Instruments P-97 puller. Prior to recording, if the cell was strongly adhered to the coverslip (as was usual 2-3 days after plating), an electrode with a blunt tip was used to scrape the surrounding area of the cell and gently maneuvered to ensure detachment of the cell from the coverslip. The “scraping” electrode was pulled from the same borosilicate capillaries used for recording electrodes. Recording pipettes were filled with a Kgluconate-based internal solution containing (in mM) 139.5 KGluconate, 1.6 MgCl2, 1 EGTA, 0.09 CaCl2, 9 HEPES,14 creatine phosphate (Tris salt), 4 mM MgATP, 0.3 mM GTP (Tris salt), pH adjusted to 7.2 with KOH. Pipette resistances were between 0.8-2 MΩ. Reported membrane potentials were corrected for a liquid junction potential of -13 mV between the internal solution and the Tyrode’s solution in which the current was zeroed before recording. Electrode tips were wrapped with thin strips of Parafilm (National Can Company) to reduce pipette capacitance and allow optimal series resistance compensation (70-80%) without oscillation. After forming a giga-ohm seal and achieving the whole-cell configuration, cells were lifted and positioned in front of a series of quartz flow pipes (250 µm ID, 350 µm OD; PolyMicro Technologies) attached with polyurethane glue to an aluminum square rod (cross section 1.5 cm x 0.5 cm) whose temperature is controlled using resistive heating elements and a feedback-controlled temperature controller (TC-344B; Warner Instruments). Cells were moved between pipes for rapid solution changes. Recordings were made with solutions at 37 °C. Cells were recorded at their natural resting membrane potential without any injection of holding current. The first action potential generated by a 1 second current injection was analyzed for the reported action potential parameters. Capsaicin sensitivity was tested at the end of the experiment by holding the cell at-70 mV in voltage clamp and briefly applying 1 µM capsaicin. Recordings were made with a dPatch amplifier (Sutter Instruments) controlled by SutterPatch software (Sutter Instruments), with currents and voltages low-pass filtered at 10 kHz using the amplifier’s built-in Bessel filter and digitized at 100 kHz or with a Multiclamp 700B amplifier (Molecular Devices) and Digidata 1322A A/D converter (Molecular Devices), controlled by Clampex 10.3.1.5 software (Molecular Devices) with currents and voltages low-pass filtered at 10 kHz using the amplifier circuitry and sampled at 100 kHz.

### Statistical Analysis

All data is presented as mean ± SEM. For statistical analysis, capacitance, resting membrane potential, rheobase, action potential property values, and capsaicin magnitude and area under the curve (AUC) values were compared using an unpaired student’s t-test. The proportion of cells with spontaneous activity and response to 20 nM capsaicin were compared using a Fisher’s exact test. Differences in number of action potentials fired in response to a ramp stimulus were compared using a non-parametric Mann-Whitney test. All data, sample size information, and statistical results can be found in Table 2. All statistical tests were performed using GraphPad Prism (Version 10.3.1).

**Table 2.** Data and Statistical Output for Electrophysiology and Calcium Imaging Experiments Conducted at UTD.

## Results

### DRGs stored in Hibernate media result in similar neuronal yield to those acutely dissociated

For experiments performed at UTD, human DRGs were either acutely dissociated or stored in Hibernate media for 4-16 hours prior to dissociation, culturing, and subsequent testing (Figure 1A). We found that both approaches resulted in similar neuronal yield per DRG (Figure 1B). We noted no impact on neuronal yield based on time in Hibernate media (Figure 1C). Similarly, yields for myeloid derived cells (CD11b+) (Figure 1D) and T cells (CD3+) (Figure 1E) using FACS on DRGs either dissociated acutely or stored in Hibernate media were comparable. Lastly, using immunocytochemistry we found that neuronal cultures from both acutely dissociated and DRGs stored in Hibernate media resulted in healthy appearing neurons with round soma and axon growth (Figure 1F).

**Figure 1.**
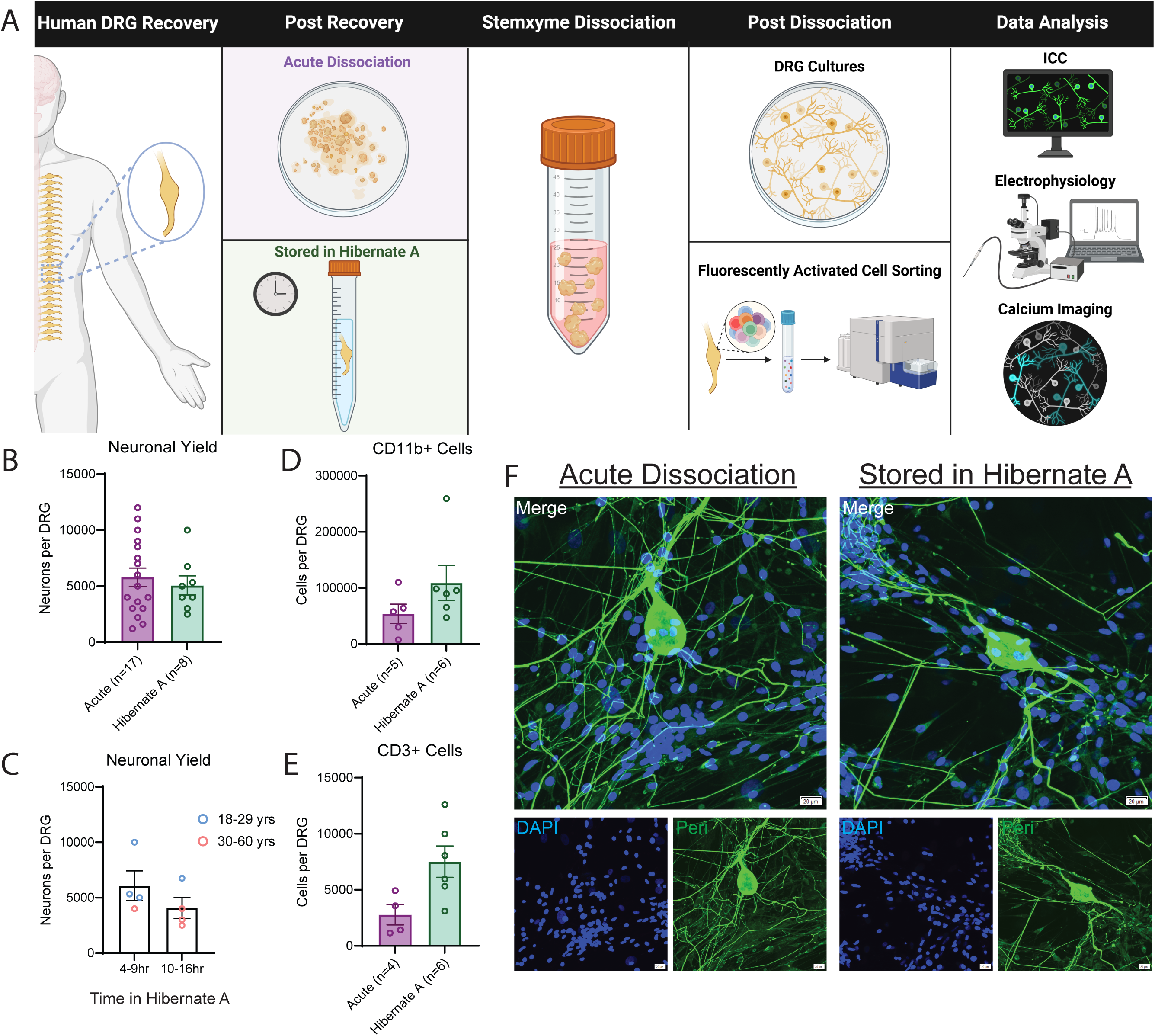
Storage in Hibernate Media Results in Similar Neuronal and Immune Cell Yield to Acutely Dissociated Tissue. A) Graphical overview of workflow for testing for differences between acutely dissociated and Hibernate media stored DRGs. B). Similar neurons per DRG were isolated regardless of whether DRG tissue was acutely dissociated or stored in Hibernate Media. C) There was a slight reduction in neurons per DRG isolated with increased time spent in Hibernate media. However, this could be driven by age of donor as older donors typically yield fewer neurons per DRG. D) There was no reduction in the number of CD11b+ myeloid cells recovered from DRG via fluorescently activated cell sorting when DRG tissue was stored in Hibernate media. E) There was no reduction in the number of CD3+ T cells recovered from DRG via fluorescently activated cell sorting when DRG tissue was stored in Hibernate media. F) Neuronal cultures from dissociated DRGs produce healthy appearing cultures with axon growth regardless of storage in Hibernate media. DRG: Dorsal Root Ganglia, ICC: Immunocytochemistry

### Acutely dissociated and Hibernate stored DRGs share similar electrophysiological properties

Next, we tested if neurons from human DRGs that were acutely dissociated or stored in Hibernate media had similar electrophysiological properties. Of the neurons tested, there was similar capacitance between the two groups (Figure 2A). Acutely dissociated neurons exhibited a modest but statistically significant elevation in resting membrane potential compared to controls (Figure 2B). There was no difference in the proportion of spontaneously active neurons, rheobase (i.e., the minimum current needed to evoke an action potential), or number of action potentials fired in response to ramp stimulus between the two groups (Figure 2C-F). When looking at the action potential properties, the only difference found was a statistical difference in the half width of the action potential with the DRGs stored in Hibernate media being slightly higher. Otherwise, there were no differences in the amplitude, rising slope, falling slope, threshold, or after hyperpolarization of the action potential (Figure 2G-M). Thus, while minor statistical differences were detected in two of the measured electrophysiological properties, the majority of the measured parameters were similar between DRG neurons acutely dissociated and those stored in Hibernate media.

**Figure 2.**
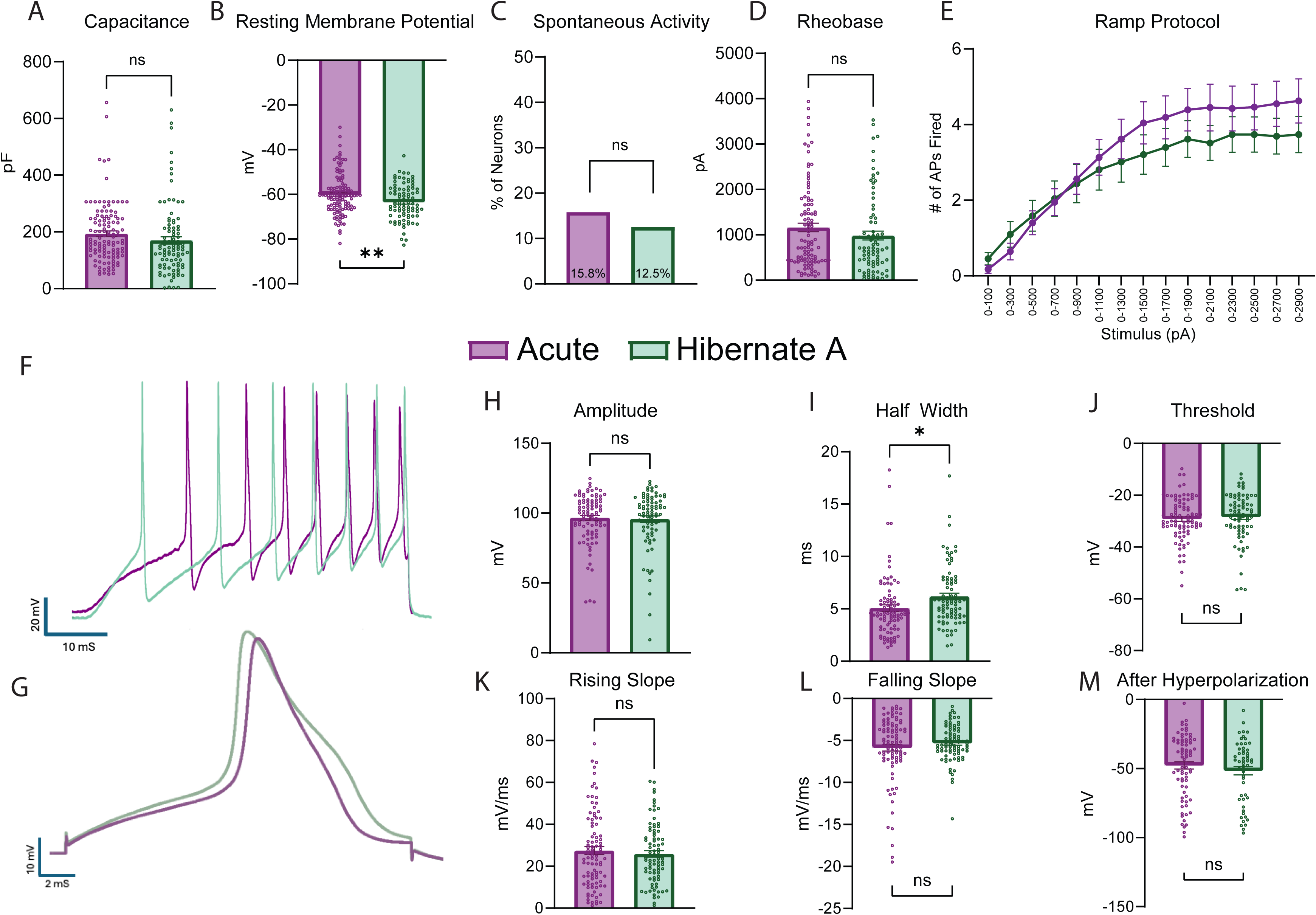
Human Neurons Isolated from DRGs Stored in Hibernate Media have Similar Electrophysiological Properties as those Acutely Dissociated. A) No difference was found in the capacitance of the neurons in each group. B) Neurons from DRGs stored in Hibernate media had a lower resting membrane potential than neurons from acutely dissociated tissue. C) There was no difference in the proportion of cells with spontaneous activity between the two groups. D) There was no difference in the rheobase between the two groups. E) There was statistical group or time effect in the number of action potentials fired during increasing stimuli between the two groups. F) Example traces of neurons from each group of the actional potentials fired during ramp stimuli. G) Example traces of neurons from each group of actional potentials in a step stimulus. H) There was no difference in the amplitude of the actional potential between each group. I) Neurons from DRGs stored in Hibernate media had a higher half width than neurons from acutely dissociated tissue. J-M) There was no difference in the threshold, rising slope, falling slope, and after hyperpolarization of the actional potential between the two groups. * p<0.05, ** p<0.01, pF: picofarads, mV: millivolt, pA: picoamperes, AP: Action Potential, ms: milliseconds

### Acutely dissociated and Hibernate media stored DRGs share similar response profiles to capsaicin stimulus

Next, we examined whether cultured neurons from human DRGs that were acutely dissociated or stored in Hibernate media had similar response profiles to capsaicin using calcium imaging. Capsaicin was used as it activates TRPV1 channels which are broadly expressed on nociceptive neurons [6; 19]. First, we found that neurons from each group had similar response rates to different concentrations of capsaicin (Figure 3A-C). We also saw a similar concentration-response effect in the magnitude of the response to increasing concentrations of capsaicin (Figure 3D). Since we had the most data for neurons treated with 20nM capsaicin, we performed a more thorough analysis of those cells. First, we showed that there was a similar proportion of cells that responded to 20nM capsaicin across several donors (Figure 3E). Of the responding neurons, there was a difference in the magnitude and area under the curve of the capsaicin response between the two groups with neurons from Hibernate media cultures being lower (Figure 3F-G). This data suggests that neurons stored in Hibernate media have similar response rates to capsaicin, but slightly lower magnitude of the capsaicin response.

**Figure 3.**
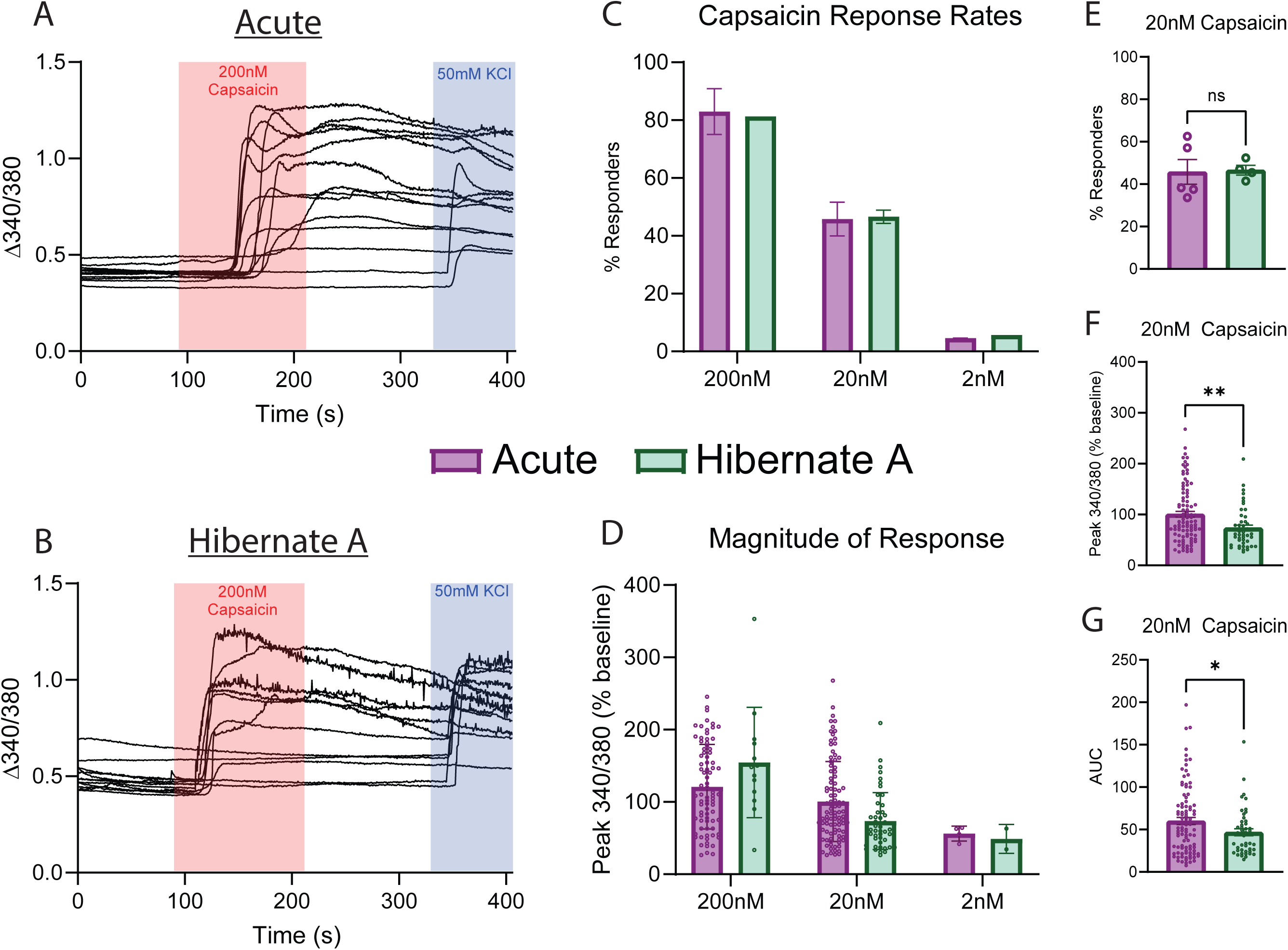
Human Neurons Isolated from DRGs Stored in Hibernate Media have Similar Calcium Image Response to Capsaicin. A) Example traces of human neurons from acutely dissociated DRG tissue challenged with 200nM capsaicin and 50mM KCl. B) Example traces of human neurons from DRG tissue stored in Hibernate media challenged with 200nM capsaicin and 50mM KCl. C) Human neurons from DRG tissue acutely dissociated or stored in Hibernate media have similar percentage response rates to descending doses of capsaicin. D) Human neurons from DRG tissue acutely dissociated or stored in Hibernate media have similar magnitude of response to descending doses of capsaicin. E) There was no difference in the proportion of neurons that responded to 20nM capsaicin between neurons from DRG tissue that was acutely dissociated or stored in Hibernate media. F) Neurons from DRGs stored in Hibernate media had a lower magnitude of response when stimulated with 20nM capsaicin when compared to neurons from DRGs acutely dissociated. G) Neurons from DRGs stored in Hibernate media had a lower area under the curve when stimulated with 20nM capsaicin when compared to neurons from DRGs acutely dissociated. * p<0.05, ** p<0.01, AUC: Area Under the Curve

### Dissociated neurons stored in Hibernate media and tested in other laboratories possess similar electrophysiological properties as neurons plated immediately after dissociation

In a separate design we acutely dissociated human DRG neurons at UTD, resuspended the cells in pre-chilled Hibernate media, stored them at 4°C, and shipped them to two different laboratories for electrophysiology testing (Figure 4A). At the University of Florida, patch clamp electrophysiology was performed with an emphasis on recording from smaller diameter neurons which is reflected in the low capacitance values (Figure 4B). The neurons had comparable electrophysiology parameters including resting membrane potential, percentage of spontaneously active neurons and action potential properties including threshold, rising slope, falling slope, half width, and after hyperpolarization when compared to neurons recorded at UTD (Figure 4C, D, G-K, Table 3). Neurons recorded at the University of Florida exhibited lower rheobase and reduced action potential amplitude compared to those recorded at UTD (Figure 4E,F, Table 3). This difference likely reflects a methodological emphasis on sampling smaller-diameter neurons at the University of Florida. [12; 20; 25].

**Figure 4.**
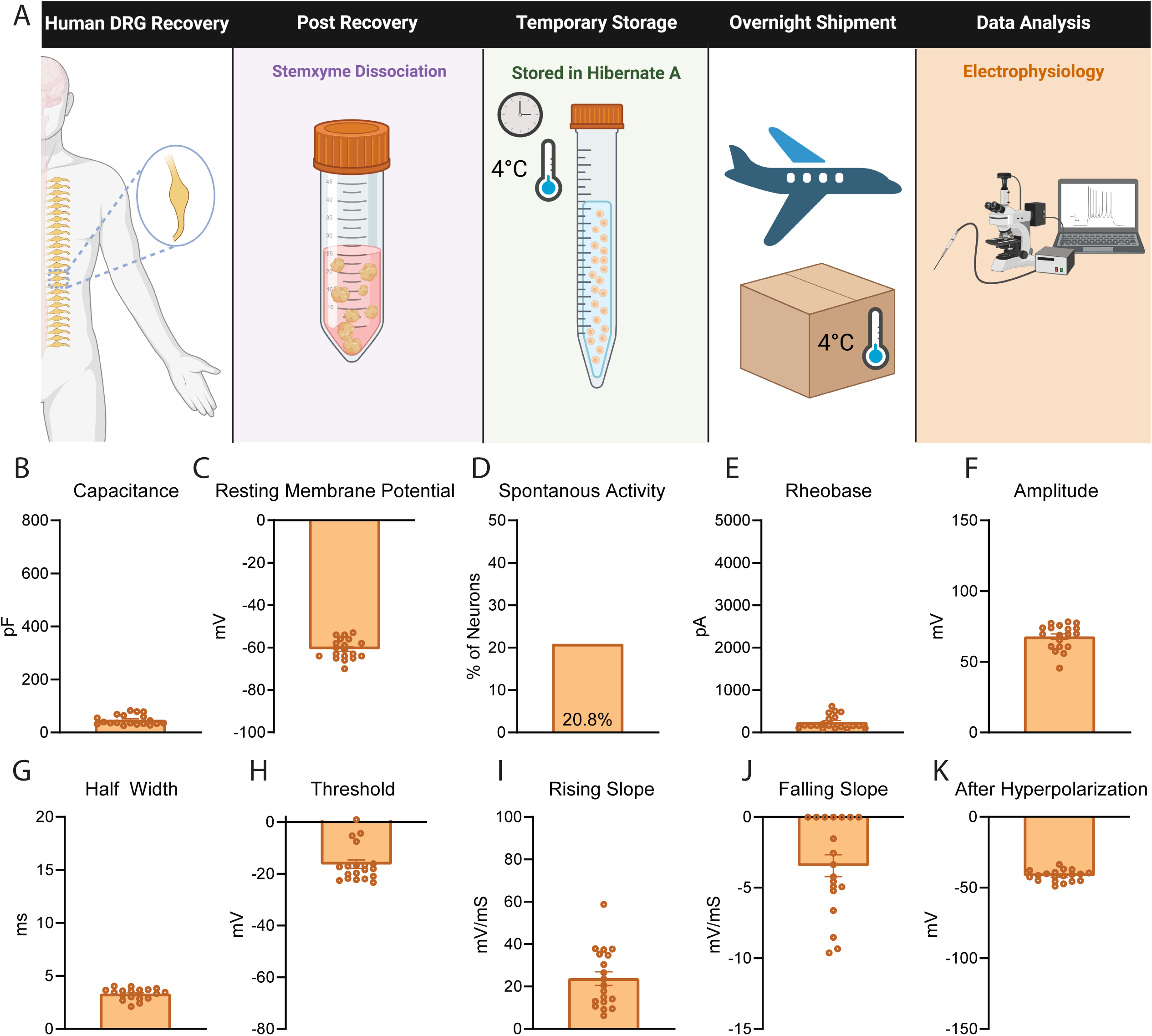
Dissociated Human DRG Neurons Stored in Hibernate Media and Recorded at University of Florida have Comparable Electrophysiological Properties to those at UTD. A) Graphical overview of workflow for testing for testing of human DRG neurons that were dissociated acutely, stored in Hibernate media, and shipped to University of Florida for testing. B) An emphasis was made to record from smaller diameter neurons at the University of Florida which is reflected in the average lower capacitance of neurons. C-D) These neurons displayed similar average resting membrane potential and percentage of spontaneously active cells to UTD neurons. E-F) These neurons exhibited a slightly lower rheobase and amplitude than UTD neurons. G-K) These neurons presented with similar half width, threshold, rising slope, falling slope, and after hyperpolarization compared to neurons measured at UTD. pF: picofarads, mV: millivolt, pA: picoamperes, ms: milliseconds

**Table 3.** Electrophysiology Data for all Experiments Conducted Across Laboratories.

Neurons were also sent to Harvard University and patch-clamp electrophysiology was performed on neurons at 37°C (Figure 5A). The neurons recorded at Harvard had similar capacitance, resting membrane potential, rheobase, amplitude, threshold, half-width, and after-hyperpolarization values when compared to neurons recorded at UTD (Figure 5B-K, Table 3). The neurons recorded at Harvard had higher rising slope and quicker falling slope when compared to neurons at UTD and UF, which can be attributed to these neurons being recorded at 37°C (Figure 5I, J, Table 3) [5; 9]. Also, 78.6% of the Harvard neurons were sensitive to high dose capsaicin (1µM) which is similar to the percentage of capsaicin responsive neurons measured via calcium imaging at UTD in both acutely dissociated neurons and in hDRGs temporarily stored in Hibernate media prior to dissociation (Figure 5L). Thus, this data demonstrates that neurons that are acutely dissociated, stored in Hibernate media, and shipped to other laboratories produce cultures of human DRG neurons with electrophysiology recordings similar to neurons that are plated and cultured immediately after dissociation.

**Figure 5.**
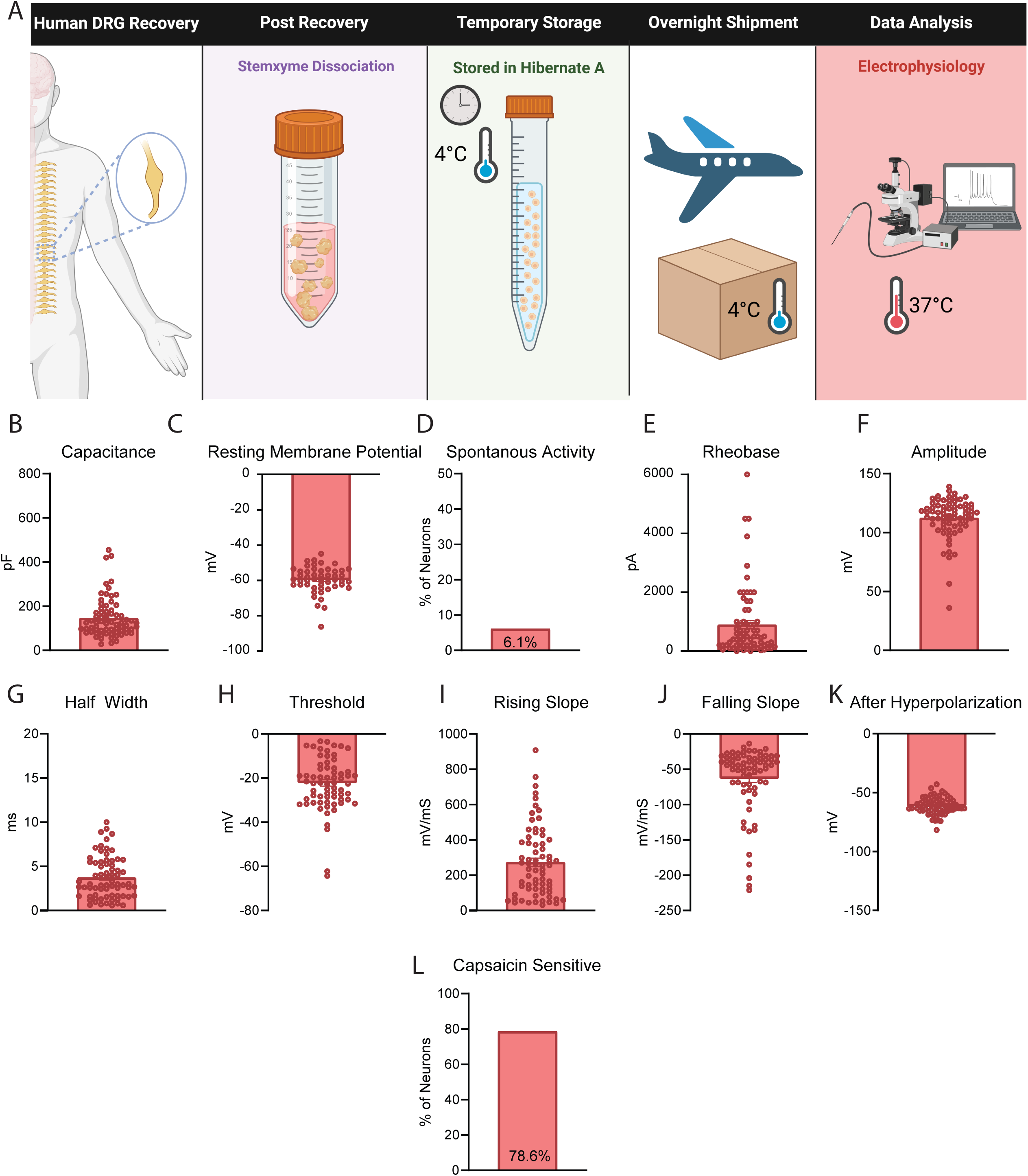
Dissociated Human DRG Neurons Stored in Hibernate Media and Recorded at 37°C at Harvard University have Comparable Electrophysiological Properties to those at UTD. A) Graphical overview of workflow for testing of human DRG neurons that were dissociated acutely, stored in Hibernate media, and shipped to Harvard University for testing at 37°C. B-H) These neurons displayed a similar capacitance, resting membrane potential, percentage of spontaneous activity, rheobase, amplitude, half width, and threshold when compared to the neurons recorded at UTD. I-J) These neurons had a steeper rising and falling slope compared to UTD measured neurons. K) These neurons displayed a similar after hyperpolarization when compared to neurons measured at UTD. L) The neurons displayed a similar percentage of capsaicin sensitivity when compared to neurons analyzed via calcium imaging at UTD. pF: picofarads, mV: millivolt, pA: picoamperes, ms: milliseconds

## Discussion

Here, we compared human DRG tissue processed in three ways: acutely dissociated, temporarily stored in Hibernate media before dissociation, and dissociated then stored in Hibernate media for shipment. Neuronal and immune cell yields were comparable across all conditions, indicating that Hibernate media preserves cell viability effectively. We also found that tissue stored in Hibernate media results in neuronal cultures that appear healthy based on axon growth and have similar electrophysiological properties and capsaicin response profiles when compared to those that were acutely dissociated. Lastly, we show that DRGs acutely dissociated, stored in Hibernate media, and shipped at 4°C to other laboratories also result in neuronal cultures with similar electrophysiological properties and capsaicin response rates when compared to neurons plated immediately after dissociation across two independent laboratories.

This work validates that human DRGs can be either temporarily stored whole or dissociated and shipped in Hibernate media to other laboratories for the use in various primary culture experiments. This agrees with prior research which demonstrated that long term (∼4 weeks) storage of rodent hippocampal tissue in Hibernate media at 4 °C can still result in viable primary neuronal cultures [4]. For shorter term storage (<16hrs), we tested the preservation of DRG tissue by keeping the DRGs fully intact in Hibernate media prior to dissociation and culturing. We found that this approach yielded very similar electrophysiological and calcium imaging data when compared to neurons from acutely dissociated DRGs. However, we found statistical differences in the resting membrane potential, half width of the action potential, and magnitude of the response to 20nM capsaicin between acutely dissociated neurons and those stored in Hibernate Media. While these differences are statistically significant, they are unlikely to be of biological significance as the magnitude of the differences is minor and could be attributed to inherent variability in organ donor cultures. One limitation in this work is we did not collect data from any donors where one DRG was dissociated acutely, and another was preserved in Hibernate media. Nevertheless, we still found similar electrophysiological and calcium imaging data across a very large sample size of neurons from many organ donors.

For longer term storage (>24hrs), we acutely dissociated DRG neurons and then resuspended the cells in Hibernate media for shipment to other laboratories. This approach allowed our group, which has extensive experience with collecting, dissociating, and culturing human DRG neurons, to share this resource with other laboratories across the country who do not have this resource or expertise. These neurons displayed very similar electrophysiological properties compared to neurons recorded in our laboratory, although some slight differences were found. At the University of Florida, an emphasis was placed on recording from smaller diameter neurons to increase probability of analysis of human nociceptors. These neurons displayed a lower average rheobase and action potential amplitude compared with neurons recorded at UTD. These differences likely stem from sampling smaller-diameter neurons, which are known to exhibit lower rheobases and reduced action potential amplitudes compared to their larger counterparts [12; 20; 25]. At Harvard, recordings were done at 37°C while recordings at UTD and University of Florida were done at room temperature. The neurons recorded at Harvard had a steeper rising and falling slope of the action potential compared to neurons recorded at UTD. These differences are likely attributable to recordings being done at 37°C which has been shown to produce quicker action potential properties due to changes in the ion channel kinetics at different temperatures [5; 9]. We have shown that acutely dissociated human DRG neurons stored in Hibernate media can be shipped to other laboratories while retaining electrophysiological properties comparable to neurons cultured immediately after dissociation.

There are several advantages to using Hibernate media when recovering DRGs from organ donors. First, temporary storage (4-16 hours) of whole DRG tissue allows the researcher to manipulate when dissociation and culturing occurs. Logistically, culturing tissue from organ donors can be challenging due to lack of control over when the collection occurs, which could be during late hours or on the weekends as hospital operating rooms are typically more available during those times. Temporary storage allows the researcher to maintain a more balanced work schedule and more flexibility in culturing while also helping to ensure appropriate completion of other already planned experiments. Also, temporary storage of tissue in Hibernate media allows for a better alignment of schedules with other researchers and scientific cores. For example, we started using Hibernate media originally to better align the DRG dissociation with our flow cytometry core for FACS at UTD which is only available during normal business hours. Secondly, long-term storage of dissociated neurons (16-42 hours) in Hibernate media allows for the sharing of this precious resource with other laboratories and universities that do not have the ability to recover this tissue on their own. Sharing human DRG neurons with other laboratories accelerates pain research, streamlines target and compound/biologic validation, and enables experiments to be completed by experts beyond the originating laboratory’s domain.

While this work demonstrates that Hibernate media can be used to store either whole DRGs or dissociated neurons for later testing, thereby helping to democratize access to viable human neural tissue, several limitations and further optimizations still need to be addressed. First, we only tested neurons from DRGs and thus do not know if this approach will be generalizable to other tissue sources. As further access and experimentation on human neural tissue progresses, Hibernate media needs to be tested on neurons from other sources such as spinal cord, trigeminal ganglia, and sympathetic chain ganglia. Another limitation is that we did not test timepoints past 16 hours for whole tissue storage, or longer than next day shipment for dissociated neuron storage. It will be important to test how long neuronal tissue can be stored in Hibernate media before large amounts of neuronal loss starts to occur. Also, longer term testing of dissociated neurons in Hibernate media will allow for more flexibility in shipping tissue to other laboratories if recoveries are on the weekend when next day shipment might not be possible or if multi-day shipments to international laboratories are required. Future work should look to test the limits of longer-term storage as well as explore the possibility of cryogenic storage which has been used in rodent and canine neuronal tissue in the past [13; 15]. A final limitation is that the majority of our experiments were tested on younger, healthy donors and thus it is unknown if this approach is translatable to all organ donors. Future work should be completed on testing Hibernate media storage solution on older donors and donors with varying medical histories such as diabetes and peripheral neuropathy.

In sum we demonstrate that storage of human DRG tissue in Hibernate media is a feasible option for producing neuronal cultures when acute dissociation is not possible. We also show that dissociation, storage in Hibernate media, and shipment to other laboratories produces viable neuronal cultures with similar electrophysiological properties as those acutely dissociated. We hope that this serves as a framework for how human neuronal tissue can be shared across various universities and laboratories who do not have access to this valuable and limited resource.

## Acknowledgements

The authors thank the organ donors and their families for their gift, members of the Southwest Transplant Alliance for supporting the tissue recovery work, and members of the Price lab for useful discussions. This research was supported by the National Institute of Neurological Disorders and Stroke of the National Institutes of Health through the PRECISION Human Pain Network (RRID:SCR_025458), part of the NIH HEAL Initiative (https://heal.nih.gov/) under award number U19NS130608 to TJP. This work was also funded by NIH grant R01NS111929 to TJP, F32NS134563 to JBL, R35-NS127216 to BPB, RF1NS131165 to RK and Department of Defense Awards HT9425-24-1-1007 and CP230147P1 to RK. The content is solely the responsibility of the authors and does not necessarily represent the official views of the National Institutes of Health.

---

**Supplemental Table 1.** Antibodies and Working Concentrations used for Fluorescently Activated Cell Sorting and Immunocytochemistry Experiments.

