## Supplemental Tables for "Enabling wider access to human molecular neuroscience research in pain: A simple preservation method for human dorsal root ganglion neurons in Hibernate A media"

### University of Texas at Dallas

| UTD Donor ID | Age | Sex | Ethnicity | COD | DRG Used | Post Recovery | Technique |
|---|---|---|---|---|---|---|---|
| UTD-DN0208 | 36 | M | White | Head Trauma/GSW | 1x Lumbar | Acute Dissociation | FACS |
| UTD-DN0240 | 19 | F | White | Anoxia/Asphyxiation/Smoke Inhalation | L4 | Acute Dissociation | FACS |
| UTD-DN0255 | 44 | M | White | CVA/Stroke | 2x Lumbar | Acute Dissociation | FACS |
| UTD-DN0269 | 29 | F | White | Head Trauma/GSW/Suicide | 2x T12 | Acute Dissociation | Ephys/$Ca^{2+}$ Imaging |
| UTD-DN0272 | 19 | M | White | Anoxia/Cardiovascular | L2,L3 | Acute Dissociation | $Ca^{2+}$ Imaging |
| UTD-DN0274 | 20 | M | White | Head Trauma/Blunt Injury/MVA | 2x L1 | Hibernate A | ICC |
| UTD-DN0278 | 35 | M | White | Anoxia/Drug Intoxication | L1,2x T4 | Acute Dissociation | ICC/Ephys/$Ca^{2+}$ Imaging |
| UTD-DN0284 | 32 | M | White | Head Trauma/MVA | L4 | Hibernate A | FACS |
| UTD-DN0285 | 23 | M | White | Anoxia/Cardiovascular/MVA | L2 | Acute Dissociation | ICC/Ephys/$Ca^{2+}$ Imaging |
| UTD-DN0286 | 53 | M | White | CVA/Stroke | L2 | Acute Dissociation | Ephys/$Ca^{2+}$ Imaging |
| UTD-DN0292 | 22 | M | White | Anoxia/Drug Intoxication | 1x Lumbar | Hibernate A | $Ca^{2+}$ Imaging |
| UTD-DN0292 | 22 | M | White | Anoxia/Drug Intoxication | 1x Lumbar | Hibernate A | FACS |
| UTD-DN0297 | 44 | F | White | Anoxia/Cardiovascular | L4 | Acute Dissociation | ICC |
| UTD-DN0298 | 46 | M | White | Anoxia/Blunt Injury/MVA | 2x L2 | Hibernate A | Ephys |
| UTD-DN0301 | 33 | M | White | Head Injury/Blunt Injury/MVA | L2 | Hibernate A | Ephys/$Ca^{2+}$ Imaging |
| UTD-DN0303 | 18 | M | Black | Head Trauma/GSW/Accident | L3 | Acute Dissociation | Ephys |
| UTD-DN0305 | 31 | M | White | Head Trauma/Accident/MVA | 2x Lumbar | Hibernate A | Ephys/$Ca^{2+}$ Imaging |
| UTD-DN0306 | 28 | F | White | Anoxia/Asphyxiation/Suicide | 2x S1 | Acute Dissociation | ICC |
| UTD-DN0315 | 21 | M | White | Head Trauma/GSW/Accident | L4 | Hibernate A | $Ca^{2+}$ Imaging |
| UTD-DN0316 | 28 | M | White | Anoxia/Drowning/Accident | 2x L5 | Acute Dissociation | Ca2+ Imaging |
| UTD-DN0318 | 20 | M | White | Head Trauma/MVA | 2x S1 | Acute Dissociation | ICC |
| UTD-DN0325 | 46 | M | White | Anoxia/Asphyxiation/Accident | 2x L2 | Hibernate A | Ephys |
| UTD-DN0327 | 22 | M | White | Head Trauma/GSW/Suicide | 2x L3 | Acute Dissociation | Ca2+ Imaging |
| UTD-DN0332 | 34 | F | Pacific Islander | Anoxia/Natural Causes | L2 | Acute Dissociation | ICC |
| UTD-DN0334 | 20 | M | White | Anoxia/Cardiovascular | T5,6,8 | Hibernate A | ICC/Ephys/$Ca^{2+}$ Imaging |
| UTD-DN0341 | 59 | M | White | Head Trauma/GSW/Suicide | T6,T12,L2,L3,L5 | Acute Dissociation | Ca2+ Imaging |
| UTD-DN0346 | 42 | F | White | Anoxia/Drug Intoxication | T10/T11/L5 | Acute Dissociation | Ca2+ Imaging |
| UTD-DN0354 | 37 | M | White | Head Trauma/Blunt Injury/non-MVA | 1x Thoracic | Acute Dissociation | Ca2+ Imaging |
| UTD-DN0354 | 37 | M | White | Head Trauma/Blunt Injury/non-MVA | L1,2,3,5 | Acute Dissociation | FACS |
| UTD-DN0356 | 19 | M | White | Head Trauma/GSW/Homicide | L2 | Acute Dissociation | Ephys |
| UTD-DN0357 | 34 | M | Black | Anoxia/Drug Intoxication | L1,L3 | Hibernate A | FACS |
| UTD-DN0358 | 31 | M | Asian | Anoxia | 2x L3 | Hibernate A | FACS |
| UTD-DN0363 | 29 | F | White | Sepsis | L1,L3 | Acute Dissociation | FACS |
| UTD-DN0367 | 45 | M | White | CVA/Stroke | 2x Lumbar | Hibernate A | FACS |
| UTD-DN0371 | 33 | M | White | Head Truama/GSW/Suicide | L2 | Acute Dissociation | Ephys |
| UTD-DN0375 | 45 | M | White | Anoxia/Cardiovascular | 2x L4 | Acute Dissociation | Ephys |
| UTD-DN0379 | 44 | F | White | CVA/Stroke | 2x L2,L5 | Acute Dissociation | Ephys |
| UTD-DN0388 | 46 | M | White | CVA/Stroke | L2,L3 | Hibernate A | FACS |

### University of Florida

| UTD Donor ID | Age | Sex | Ethnicity | COD | DRG Used | Post Recovery | Technique |
|---|---|---|---|---|---|---|---|
| UTD-DN0322 | 60 | M | White | Head Trauma/Blunt Injury/MVA | 2x T12 | Dissociated Neurons in Hibernate A | Ephys |
| UTD-DN0326 | 39 | F | White | Anoxia/Asphyxiation | L2 | Dissociated Neurons in Hibernate A | Ephys |
| UTD-DN0330 | 41 | M | White | CVA/Stroke | 1x Thoracic | Dissociated Neurons in Hibernate A | Ephys |
| UTD-DN0334 | 20 | M | White | Anoxia/Cardiovascular | T7 | Dissociated Neurons in Hibernate A | Ephys |

### Harvard University

| UTD Donor ID | Age | Sex | Ethnicity | COD | DRG Used | Post Recovery | Technique |
|---|---|---|---|---|---|---|---|
| UTD-DN0397 | 58 | F | White | Anoxia/Cardiovasular | 2x T11 | Dissociated Neurons in Hibernate A | Ephys |
| UTD-DN0399 | 24 | F | White | Cardiovascular/Overdose | 1x Lumbar | Dissociated Neurons in Hibernate A | Ephys |
| UTD-DN0408 | 55 | M | | Anoxia/Cardiovasular | 2x T12 | Dissociated Neurons in Hibernate A | Ephys |
| UTD-DN0411 | 25 | F | | Anoxia/Cardiovasular | L2,L3 | Dissociated Neurons in Hibernate A | Ephys |
| UTD-DN0418 | 2 | M | | Drowning | L3,L4 | Dissociated Neurons in Hibernate A | Ephys |

| **Electrophysiology** | | | | |
|---|---|---|---|---|
| Capacitance | | | | |
| **Group** | **Mean** | **SD** | **n** | **t-test** |
| Acute | 193.95 | 104.16 | 118 | $t_{209}$=1.50, p=0.13 |
| Hibernate A | 170.12 | 126.20 | 93 | |
| Resting Membrane Potential | | | | |
| **Group** | **Mean** | **SD** | **n** | **t-test** |
| Acute | -60.11 | 9.09 | 114 | $t_{199}$=2.77, p<0.01 |
| Hibernate A | -63.50 | 7.90 | 87 | |
| Spontaneous Activity | | | | |
| **Group** | **% Responders** | | **n** | **Fishers Test** |
| Acute | 15.83% | | 120 | p=0.56 |
| Hibernate A | 12.50% | | 96 | |
| Rheobase | | | | |
| **Group** | **Mean** | **SD** | **n** | **t-test** |
| Acute | 1161.29 | 941.78 | 101 | $t_{181}$=1.32, p=0.19 |
| Hibernate A | 982.32 | 881.06 | 82 | |
| Ramp | | | | |
| **Group** | **Mean** | **SD** | **n** | **Mann-Whitney** |
| Acute | 3.24 | 1.53 | 15 | U=76, p=0.14 |
| Hibernate A | 2.81 | 1.06 | 15 | |
| Amplitude | | | | |
| **Group** | **Mean** | **SD** | **n** | **t-test** |
| Acute | 96.54 | 17.68 | 94 | $t_{174}$=0.28, p=0.78 |
| Hibernate A | 95.73 | 20.69 | 82 | |
| Half Width | | | | |
| **Group** | **Mean** | **SD** | **n** | **t-test** |
| Acute | 5.06 | 2.94 | 92 | $t_{172}$=2.50, p=0.01 |
| Hibernate A | 6.18 | 2.90 | 82 | |
| Threshold | | | | |
| **Group** | **Mean** | **SD** | **n** | **t-test** |
| Acute | -29.15 | 8.18 | 89 | $t_{167}$=0.55, p=0.58 |
| Hibernate A | -28.41 | 9.23 | 80 | |
| Rising Slope | | | | |
| **Group** | **Mean** | **SD** | **n** | **t-test** |
| Acute | 27.41 | 18.06 | 94 | $t_{174}$=0.60, p=0.55 |
| Hibernate A | 25.91 | 14.39 | 82 | |
| Falling Slope | | | | |
| **Group** | **Mean** | **SD** | **n** | **t-test** |
| Acute | -5.87 | 3.85 | 94 | $t_{174}$=1.09, p=0.28 |
| Hibernate A | -5.34 | 2.17 | 82 | |
| After Hyperpolarization | | | | |
| **Group** | **Mean** | **SD** | **n** | **t-test** |
| Acute | -47.69 | 22.82 | 77 | $t_{130}$=0.96, p=0.34 |
| Hibernate A | -51.54 | 22.45 | 55 | |
| **Calcium Imaging** | | | | |
| 20nM Capsaicin % Responders | | | | |
| **Group** | **% Responders** | | **n** | **Fishers Test** |
| Acute | 41.38% | | 232 | p=0.35 |
| Hibernate A | 45.45% | | 101 | |
| 20nM Capsaicin Magnitude of Response | | | | |
| **Group** | **Mean** | **SD** | **n** | **t-test** |
| Acute | 100.43 | 55.42 | 96 | $t_{140}$=2.96, p<0.01 |
| Hibernate A | 73.47 | 39.46 | 46 | |
| 20nM Capsaicin AUC | | | | |
| **Group** | **Mean** | **SD** | **n** | **t-test** |
| Acute | 60.05 | 40.63 | 96 | $t_{140}$=1.99, p=0.04 |
| Hibernate A | 46.79 | 27.79 | 46 | |

| | UTD Acute | | | UTD Hibernate A | | | University of Florida Hibernate A | | | Harvard University Hibernate A | | |
|---|---|---|---|---|---|---|---|---|---|---|---|---|
| Measures | Mean | SD | n | Mean | SD | n | Mean | SD | n | Mean | SD | n |
| Capacitance | 193.95 | 104.16 | 118 | 170.12 | 126.20 | 93 | 47.20 | 19.07 | 19 | 146.29 | 89.31 | 70 |
| Resting Membrane Potential | -60.11 | 9.09 | 114 | -63.50 | 7.90 | 87 | -60.68 | 4.75 | 19 | -59.57 | 7.53 | 49 |
| Rheobase | 1161.29 | 941.78 | 101 | 982.32 | 881.06 | 82 | 240.96 | 165.60 | 19 | 896.00 | 1177.19 | 70 |
| Amplitude | 96.54 | 17.68 | 94 | 95.73 | 20.69 | 82 | 67.78 | 8.85 | 19 | 112.49 | 17.36 | 70 |
| Half Width | 5.06 | 2.94 | 92 | 6.18 | 2.90 | 82 | 3.30 | 0.53 | 19 | 3.72 | 2.37 | 70 |
| Threshold | -29.15 | 8.18 | 89 | -28.41 | 9.23 | 80 | -16.30 | 7.01 | 19 | -22.12 | 12.64 | 69 |
| Rising Slope | 27.41 | 18.06 | 94 | 25.91 | 14.39 | 82 | 23.75 | 13.86 | 19 | 272.80 | 199.19 | 70 |
| Falling Slope | -5.87 | 3.85 | 94 | -5.34 | 2.17 | 82 | -3.44 | 3.78 | 19 | -62.92 | 48.09 | 70 |
| After Hyperpolarization | -47.69 | 22.82 | 77 | -51.54 | 22.45 | 55 | -41.70 | 3.92 | 19 | -61.36 | 6.96 | 70 |

| Fluorescently Activated Cell Sorting | | | |
|---|---|---|---|
| **Antibody** | **Company** | **Product #** | **Dilution/Volume** |
| Myelin Antibodies | Miltenyi | 130-096-733 | 1:10 |
| Zombie UV Fixable Dye | Biolegend | 423107 | 1:100 |
| Human TruStain FcX | Biolegend | 422302 | 5µL |
| Anti-CD45 (BV605) | Biolegend | 304042 | 5µL |
| Anti-CD11b (PECy7) | Biolegend | 301322 | 5µL |
| Anti-CD3 (APC) | Biolegend | 300411 | 5µL |
| Immunocytochemistry | | | |
| **Antibody** | **Company** | **Product #** | **Dilution** |
| Peripherin | ENCOR | CPCA-Peri | 1:1000 |
| Goat Anti-Chicken AF 488 | Invitrogen | A11039 | 1:2000 |
| DAPI | Cayman Chemical | 14285 | 1:5000 |
